# Bulk RNA-seq data of an Adherent-Invasive *Escherichia coli* Strain ZVL and Its *exuR*-Knockout Mutant

**DOI:** 10.64898/2026.11.24.768809

**Authors:** Maria Mikhailycheva, Nikita Trusov, Maria Galyamina, Dmitriy Shirokov, Diana Urazaeva, Aleksandr Avshalumov, Natalia Borzenko, Andrei Semenov, Olga Pobeguts, Alexey Gorbachev

## Abstract

Adherent-invasive Escherichia coli (AIEC) is a subtype of pathogenic E. coli associated with Crohn’s disease. AIEC strains are characterized by the ability to adhere to and invade epithelial cells, survive and replicate within macrophages. Despite many years of research, the mechanisms of virulence and pathogenicity still remain not fully understood. Previous studies have shown that the virulence-associated phenotype of some isolates can be reversibly eliminated by changing the growth medium, specifically by switching from M9 minimal medium supplemented with propionate to M9 supplemented with glucose. This phenomenon was also observed in ZVL, an AIEC isolate from our laboratory collection. Since certain sugar acids, particularly hexuronates, have been shown to contribute to gut colonization by other bacterial pathogens, their role in AIEC has not been reported in the literature. Therefore, we aimed to investigate whether the hexuronate degradation pathway influences the adhesion and invasion properties of AIEC. To this end, we generated a CRISPR-based deletion mutant lacking exuR, which encodes a transcriptional repressor of the Ashwell pathway, and performed bulk RNA sequencing of wild-type ZVL and the ΔexuR mutant. Both strains were grown in M9 minimal medium supplemented with either glucose or sodium propionate. Caco-2 cell assays were used to assess bacterial adhesion and invasion under both growth conditions. Our results suggest that deletion of *exuR* affects the invasion phenotype following growth in propionate-supplemented medium.

**Dataset:** https://www.ncbi.nlm.nih.gov/bioproject/1542820.

## 1. Introduction

Crohn’s disease is a chronic, relapsing inflammatory bowel disease characterized by transmural inflammation that can affect any part of the gastrointestinal tract and may be accompanied by extraintestinal manifestations [1]. Adherent-invasive *Escherichia coli* (AIEC), associated with Crohn’s disease, may contribute to intestinal inflammation through its ability to adhere to and invade intestinal epithelial cells and evade immune clearance by surviving and replicating within macrophages. Understanding how these bacteria develop a virulent phenotype may help clarify their role in disease and find a potential target for new therapeutic approaches. Despite extensive research, no common mechanism explaining virulence across all AIEC strains has yet been established [2].

The ability of some AIEC isolates to change their phenotype in response to different growth media also remains unexplained. Previous experiments showed that bacteria grown in minimal medium containing either propionate or glucose as the sole carbon source exhibited differences in adhesion to and invasion of intestinal epithelial cells. Propionate-grown bacteria demonstrated increased adhesion and invasion compared with glucose-grown bacteria, whereas subsequent cultivation in glucose-containing medium reversed these properties [3]. More efficient intestinal colonization was also observed in a mouse model following growth with propionate [4]. Because propionate is a common short-chain fatty acid in the gut and has antibacterial properties, this ability of AIEC to respond to propionate may help them occupy an intestinal niche and gain a competitive advantage. In addition to experiments conducted by another group with the model strain LF82, we also demonstrated this response in ZVL, an isolate from our laboratory collection [3]. We subsequently used these growth conditions as experimental models to reproduce virulent and non-virulent phenotypes within the same bacterial strain, thereby allowing us to investigate phenotypic differences independently of genetic background.

In preliminary unpublished experiments with ZVL, we observed substantial changes in the expression of the *uxu* and *uxa* gene clusters and the transcriptional regulator gene *exuR* when cultures were switched between propionate- and glucose-containing media. These clusters encode enzymes involved in the catabolism of glucuronate and galacturonate. Although hexuronate utilization can promote intestinal colonization by other enteric pathogens [5], the role of this pathway and its regulators, ExuR and UxuR, in AIEC host-cell interactions has not been reported. Hexuronate-dependent regulation may extend beyond catabolism: in non-pathogenic laboratory strain *E. coli* K-12 MG1655, in which D-galacturonate and D-glucuronate had distinct effects on the expression of genes involved in adhesion, flagellar function, and biofilm regulation [6].

ExuR is a transcriptional regulator that controls hexuronate metabolism (Ashwell pathway). In enterohaemorrhagic *E. coli* (EHEC), ExuR has been shown to regulate virulence gene expression by directly binding to the regulatory region of *ler* and activating its transcription in the absence of galacturonic acid. Ler, in turn, activates genes within the locus of enterocyte effacement (LEE), which encodes the type III secretion system (T3SS) and other factors required for attaching-and-effacing lesion formation [7]. These findings link hexuronate metabolism to virulence regulation in EHEC and provide a rationale for selecting *exuR* for deletion in ZVL, although they do not establish a common equivalent regulatory mechanism in AIEC.

## 2. Results

### 2.1. Experimental Design and Associated Phenotypes

*E. coli* strain ZVL is a clinical isolate previously recovered from a patient diagnosed with Crohn’s disease and exhibits adherent-invasive properties [3]. Previous *in silico* analysis of the ZVL genome assembly with ECTyper predicted serotype O12:H20, and analysis with EzClermont assigned ZVL to phylogroup A. A Δ*exuR* mutant was derived from ZVL by targeted gene deletion using CRISPR-Cas9-mediated homologous recombination, and the deletion was confirmed by Sanger sequencing and whole-genome sequencing (WGS). Transcriptomes were obtained for both wild-type ZVL and the mutant grown in M9 minimal medium supplemented with glucose or sodium propionate, representing four strain-medium combinations, with three replicates per condition. The final dataset contains 12 samples for subsequent differential-expression analysis (Tables 2 and 3).

**Table 1.**
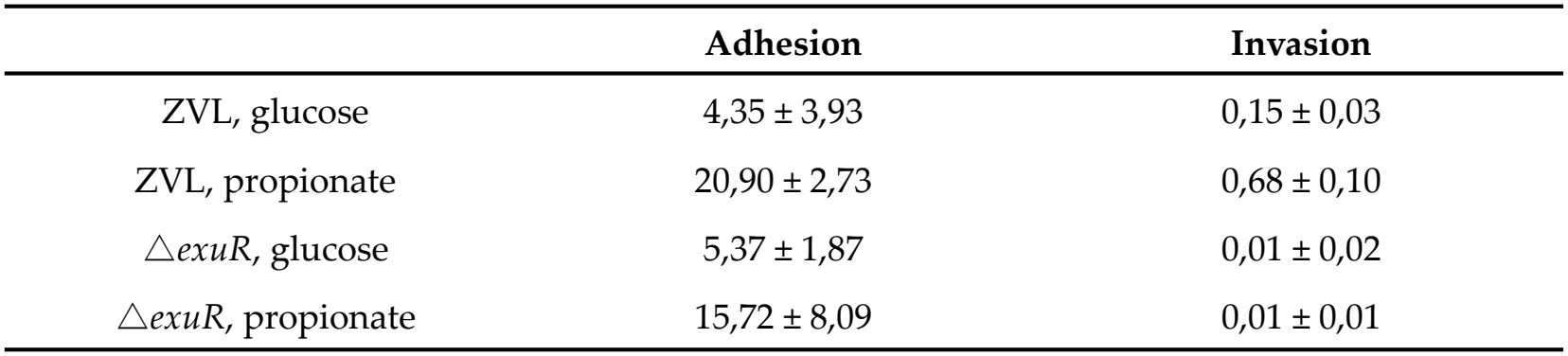
Adhesion and invasion phenotypes in Caco-2 cells assessed using the gentamicin protection assay.

|  | **Adhesion** | **Invasion** |
|---|---|---|
| ZVL, glucose | 4,35 ± 3,93 | 0,15 ± 0,03 |
| ZVL, propionate | 20,90 ± 2,73 | 0,68 ± 0,10 |
| △*exuR*, glucose | 5,37 ± 1,87 | 0,01 ± 0,02 |
| △*exuR*, propionate | 15,72 ± 8,09 | 0,01 ± 0,01 |

**Table 2.** Sample identifiers, sequencing runs, read-pair counts, and repository accessions. The suffixes g and p indicate M9 medium supplemented with glucose and propionate, respectively. The prefixes Z and E indicate wild-type ZVL and the Δ*exuR* mutant, respectively. Numbers 1-3 identify replicates.

| Run | Sample | Paired Reads (M) | BioSample | SRA |
|---|---|---|---|---|
| V350316360 | Zg1 | 22,1 | SAMN64159954 | SRR41147851 |
| V350316360 | Zg2 | 20,8 | SAMN64159955 | SRR41147850 |
| V350296481 | Zg3 | 23,2 | SAMN64159956 | SRR41147847 |
| V350316360 | Zp1 | 25,5 | SAMN64159957 | SRR41147846 |
| V350316360 | Zp2 | 22,1 | SAMN64159958 | SRR41147845 |
| V350296481 | Zp3 | 22,3 | SAMN64159959 | SRR41147844 |
| V350316360 | Eg1 | 24,9 | SAMN64159960 | SRR41147843 |
| V350316360 | Eg2 | 27,2 | SAMN64159961 | SRR41147842 |
| V350296481 | Eg3 | 21,7 | SAMN64159962 | SRR41147841 |
| V350316360 | Ep1 | 25,2 | SAMN64159963 | SRR41147840 |
| V350316360 | Ep2 | 23,9 | SAMN64159964 | SRR41147849 |
| V350296481 | Ep3 | 24,7 | SAMN64159965 | SRR41147848 |

**Table 3.** Quality statistics following STAR alignment to the ZVL genome assembly. The 5’-3’ bias metric describes the relative coverage at the beginning and end of transcripts.

| Sample | % rRNA | Duplication | 5'-3' bias | Aligned reads | Uniq aligned |
|---|---|---|---|---|---|
| Eg1 | 0.0 | 53.4% | 1.12 | 97.5% | 83.7% |
| Eg2 | 0.0 | 54.3% | 1.12 | 97.2% | 83.0% |
| Eg3 | 0.0 | 57.8% | 1.21 | 81.5% | 73.3% |
| Ep1 | 0.0 | 64.0% | 0.90 | 96.3% | 77.0% |
| Ep2 | 0.0 | 65.3% | 0.86 | 95.7% | 76.0% |
| Ep3 | 0.0 | 67.5% | 0.87 | 82.7% | 71.4% |
| Zg1 | 0.0 | 50.6% | 1.08 | 93.3% | 81.9% |
| Zg2 | 0.0 | 48.8% | 1.09 | 97.1% | 84.6% |
| Zg3 | 0.0 | 62.9% | 1.13 | 81.5% | 73.8% |
| Zp1 | 0.0 | 64.7% | 1.01 | 93.6% | 70.3% |
| Zp2 | 0.0 | 63.6% | 0.99 | 95.6% | 70.5% |
| Zp3 | 0.0 | 70.8% | 1.11 | 69.7% | 56.3% |

**Table 4.**
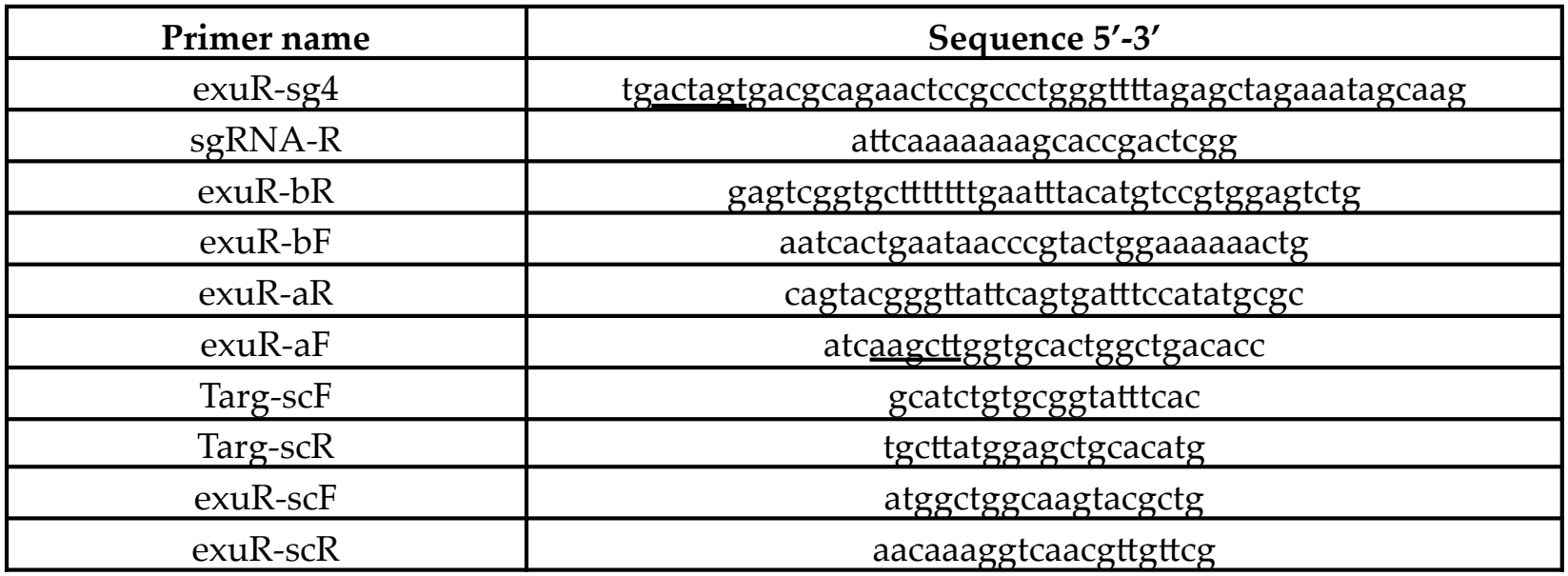
Primers used for genome editing (restriction sites are underlined).

The adhesion and invasion phenotypes were assessed using Caco-2 cells, a human colorectal adenocarcinoma cell line widely used as an *in vitro* model of the intestinal epithelial barrier [8]. The phenotypes are summarized in Table 1 and Figure 1. Mean adhesion to Caco-2 cells was higher after growth in propionate-supplemented M9 medium than after growth in glucose-supplemented M9 medium for both wild-type ZVL and the Δ*exuR* mutant. Mean invasion was also higher after growth with propionate in wild-type ZVL, whereas it remained at 0.01% in the mutant under both conditions, below the 0.1% threshold for classifying strains as invasive reported in the literature [2]. The assay reported as adhesion measured total cell-associated bacteria, including both attached and internalized organisms, whereas the gentamicin protection assay quantified viable intracellular bacteria recovered after antibiotic treatment. The latter readout reflects bacterial entry and subsequent intracellular survival over the assay period rather than entry alone. The observed pattern is therefore consistent with a contribution of *exuR* to the propionate-associated host-cell interaction phenotype, but does not demonstrate direct regulation of invasion by ExuR.

**Figure 1.**
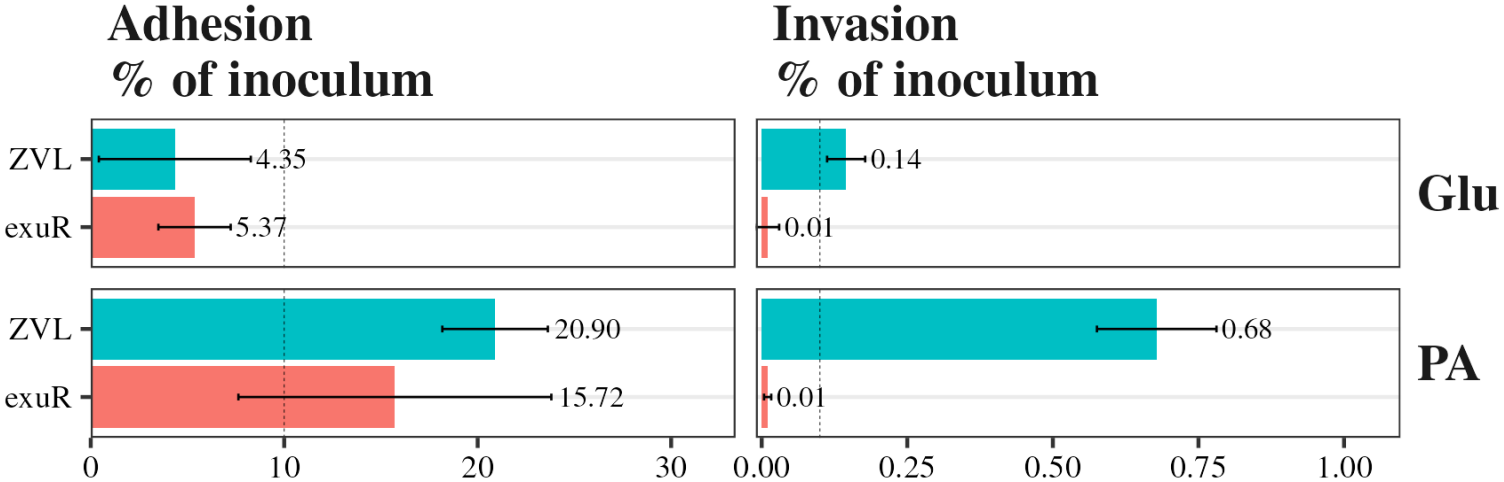
Phenotypic characterization of wild-type ZVL and Δ*exuR*-ZVL strains after five passages on minimal M9 medium supplemented with glucose (Glu) or propionate (PA). Dashed lines indicate the thresholds used to distinguish adherent from nonadherent phenotypes and invasive from noninvasive phenotypes.

### 2.2. Sequencing Records and Metadata

Raw paired-end sequencing data are identified in SRA, with linked sample descriptions in BioSample and project-level information in BioProject PRJNA1542820. The twelve records contain approximately 283.6 million read pairs in total, equivalent to approximately 56.7 Gb of sequence at 2 × 100 bp (Table 2). SRA records can be obtained through repository download services and exported as paired FASTQ files for reanalysis.

Sequencing was performed in two runs. Two replicates from each condition were sequenced in run V350316360, and the third was sequenced in run V350296481 (Table 2). Because each sequencing run included samples from all four strain-medium combinations, the sequencing run was not confounded with either strain or medium. Nevertheless, all four samples from the second run had lower alignment rates (Table 3), indicating a run-associated difference that should be considered in downstream analysis.

### 2.3. Alignment Quality and Sample Similarity

Table 3 summarizes the alignment and quality metrics from the MultiQC report generated by the Nextflow nf-core/rnaseq pipeline [9]. The complete genome assembly of strain ZVL and its corresponding annotation were used for quality control and alignment. They are not yet publicly available, which currently limits exact reproduction of the reported alignment and coverage results. Until the ZVL assembly is published, the standard *E. coli* K-12 MG1655 reference genome and its corresponding annotation can be used for reanalysis.

Total alignment rates ranged from 69.7% to 97.5%, and uniquely aligned reads ranged from 56.3% to 84.6%. Duplication estimates ranged from 48.8% to 70.8%. Residual rRNA was reported as 0.0% for every record at the displayed precision. This should not be interpreted as proof that all ribosomal sequences were absent. Following STAR alignment, the sample-distance heatmap showed clustering by experimental condition, with additional differences between sequencing runs (Figure 2; Table 2).

**Figure 2.**
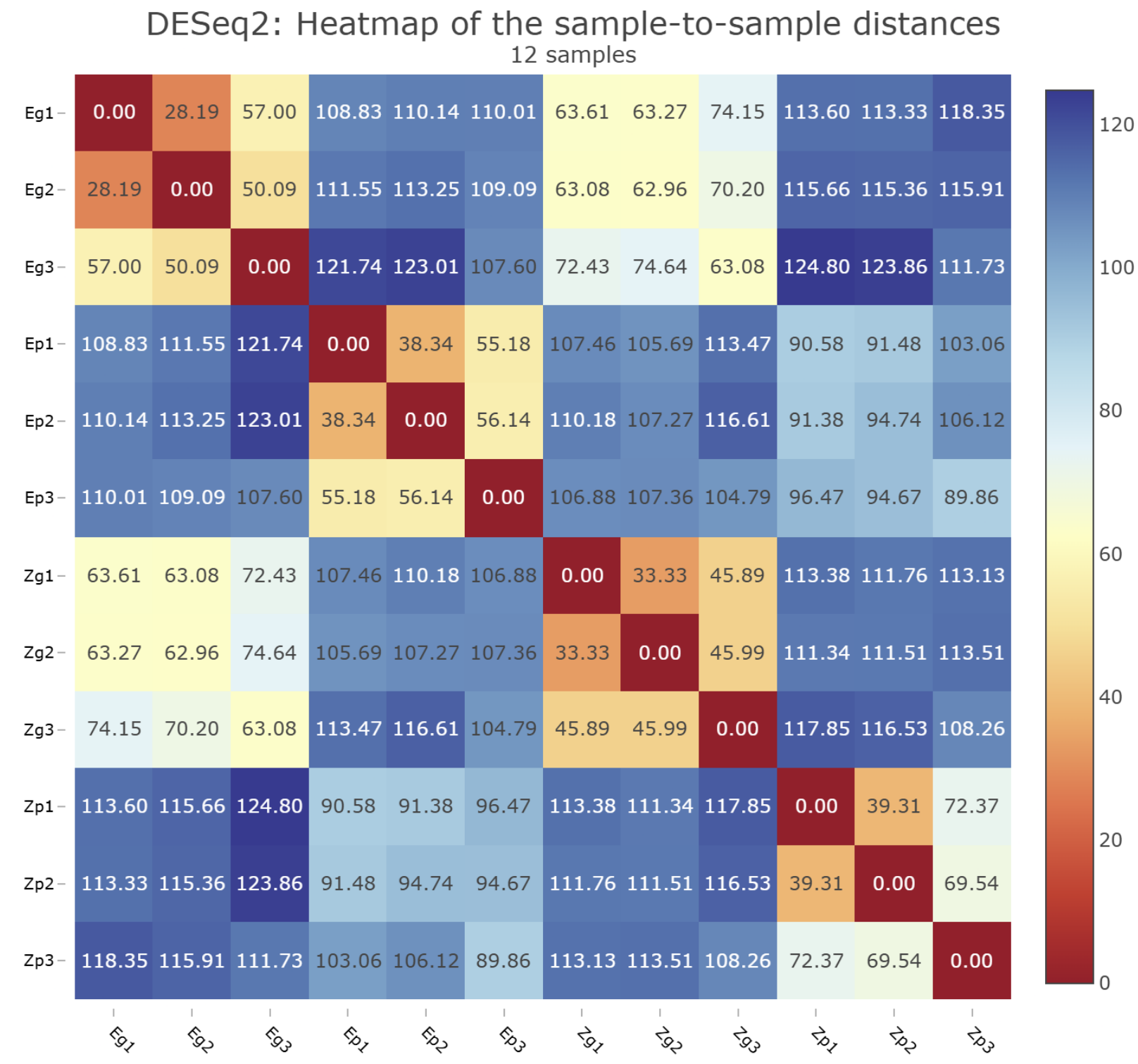
A clustered heatmap showing pairwise Euclidean distances between the samples after read count normalisation with the VST transformation.

Principal component analysis (PCA) of read counts after variance-stabilizing transformation with the DESeq2 [10] function separated samples grown with glucose from those grown with propionate along PC1 (Figure 3). PC2 separated wild-type ZVL from the Δ*exuR* mutant more clearly in propionate than in glucose.

**Figure 3.**
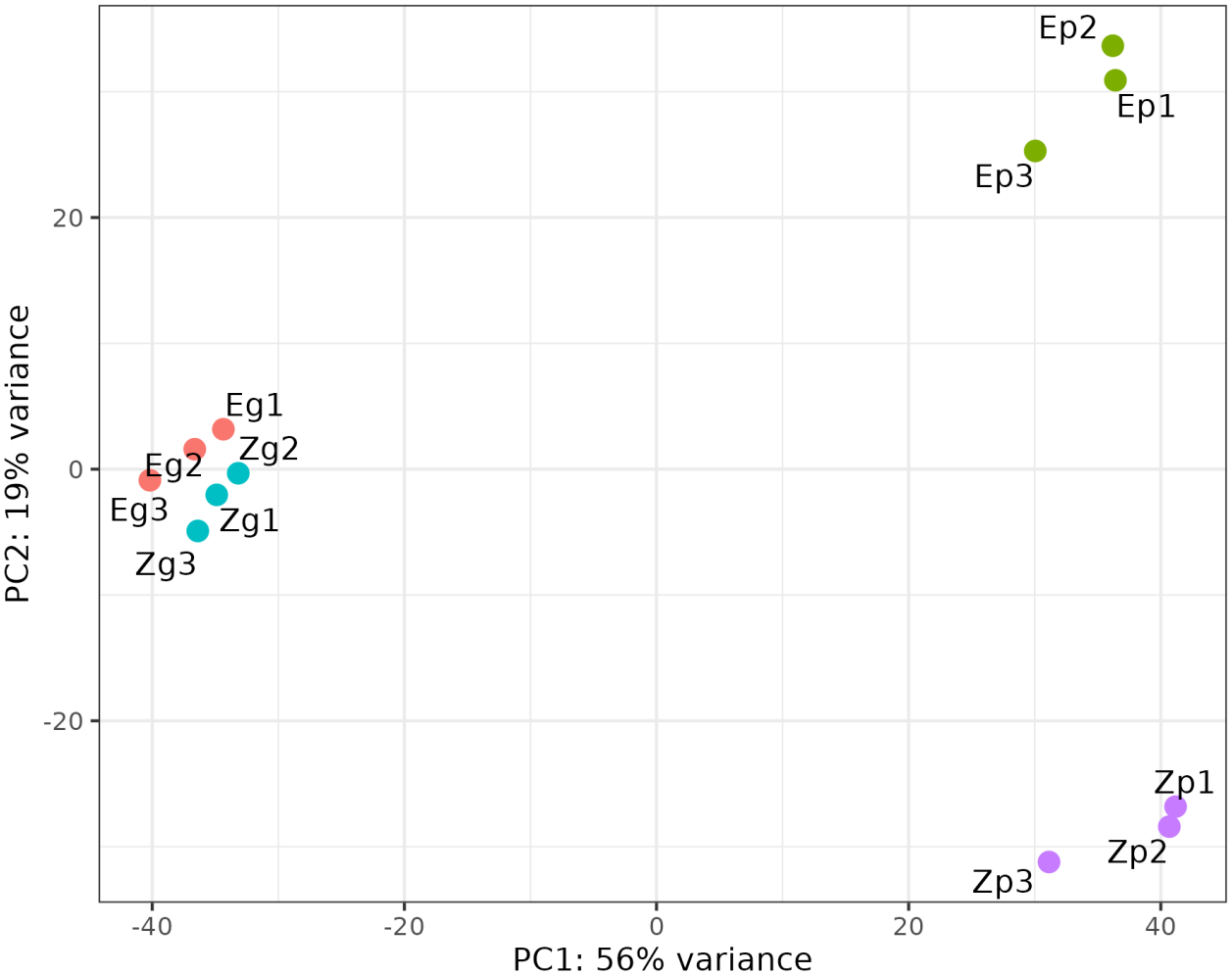
Principal component analysis of the 12 RNA-seq samples. Each point represents one replicate. Colors indicate the four strain-medium combinations. Z and E denote wild-type ZVL and the Δ*exuR* mutant, respectively. The letters g and p denote glucose and propionate, respectively, and the numbers 1-3 identify replicates.

We performed differential expression analysis using DESeq2 and found 567 upregulated and 543 downregulated genes in the Δ*exuR* mutant relative to wild-type ZVL when both strains were grown on propionate (the adjusted p-value < 0.01 and the absolute log_2_FoldChange > 1). In wild-type ZVL, 767 genes were upregulated and 695 were downregulated during growth on propionate compared with glucose. For each contrast, the upregulated and downregulated gene sets were analyzed separately for enrichment of Gene Ontology (GO) Biological Process (BP) terms using genome annotation generated by Bakta [11].

Genes upregulated in wild-type ZVL grown in propionate-supplemented medium compared with glucose-supplemented medium, as well as genes upregulated in wild-type ZVL relative to the Δ*exuR* mutant during growth on propionate, shared several GO terms. They included terms associated with bacterial motility, such as flagellar assembly and chemotaxis, thiamine biosynthesis, which supports pyruvate metabolism, and arginine catabolism (Figure 4). Flagella are known to play important roles in bacterial adhesion and invasion by promoting motility and regulating type I pilus expression [12]. Another enriched GO term was protein secretion via the type III secretion system (T3SS) (GO:0030254). However, the genes contributing to this enrichment belonged to the *fli* operon rather than encoding components of a canonical T3SS. This apparent association is explained by sequence homology between flagellar export apparatus components and the protein secretion machinery of the T3SS. Similarly, genes downregulated in ZVL relative to the Δ*exuR* mutant during growth on propionate and genes downregulated in ZVL grown on propionate compared with glucose shared an enriched term of the enterobactin biosynthetic process. Siderophores are generally considered to promote bacterial virulence and gut colonization [13]; however, our results suggest an opposite effect in this case. This may indicate that other factors outweigh the contribution of siderophores, or that excessive siderophore production could negatively affect eukaryotic cell viability. Overall, in some cases, the effect of *exuR* deletion seems to mimic the effect of shifting bacteria from virulence-inducing to non-inducing growth conditions.

**Figure 4.**
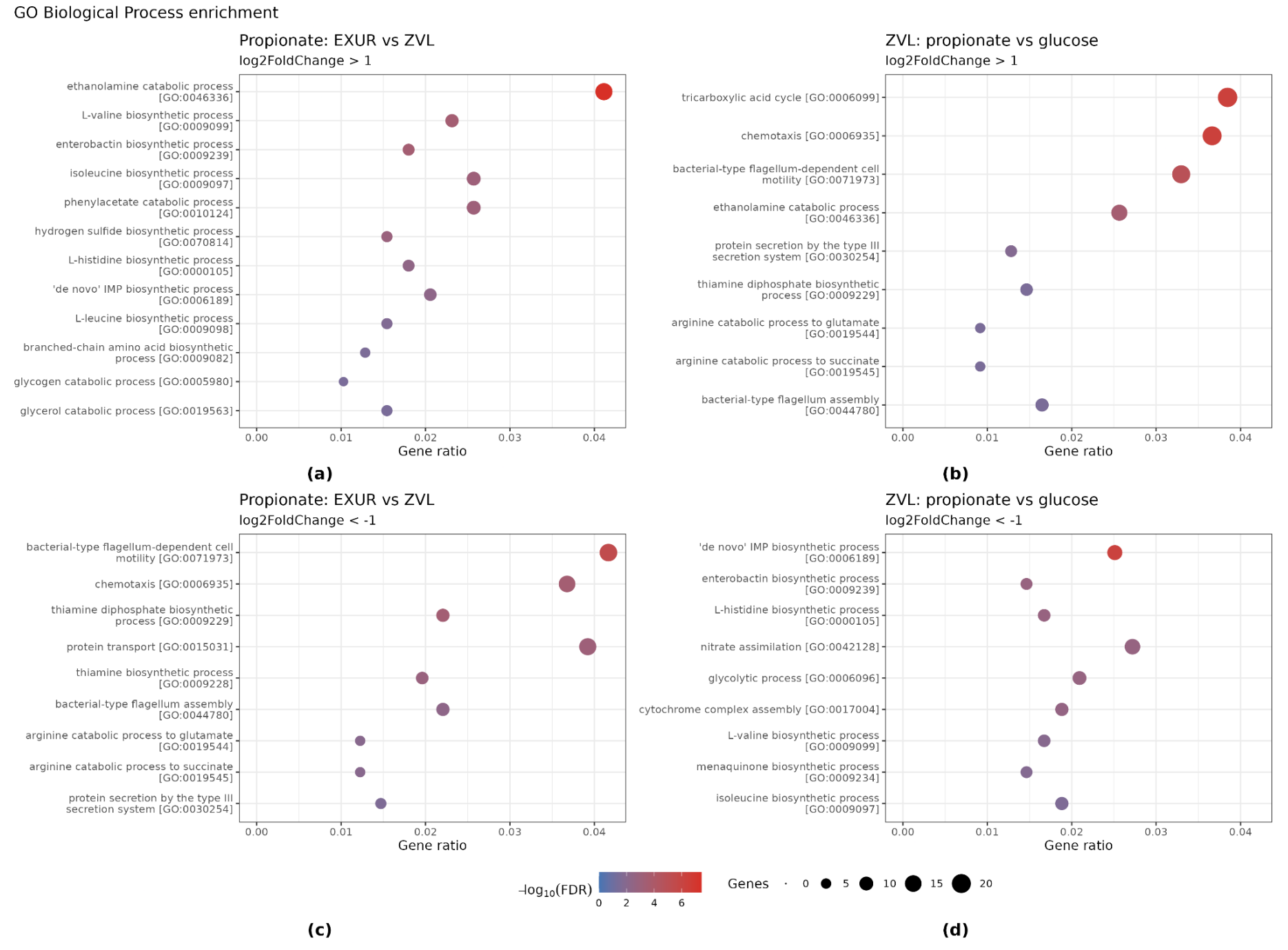
Gene Ontology Biological Process (GO BP) enrichment among differentially expressed genes. All gene sets were selected using an adjusted p-value < 0.01. All GO BP terms with a Benjamini-Hochberg FDR < 0.05 are shown. The Gene ratio is the fraction of selected genes with a BP annotation that are annotated to the indicated term. Point size represents the number of selected genes annotated to the term, and color represents -log_10_(FDR). (**a**) Δ*exuR* versus ZVL during growth on propionate, enrichment among upregulated genes (log_2_FoldChange > 1). (**b**) Propionate versus glucose in ZVL, enrichment among upregulated genes (log_2_FoldChange > 1). (**c**) Δ*exuR* versus ZVL during growth on propionate, enrichment among downregulated genes (log_2_FoldChange < -1). (**d**) Propionate versus glucose in ZVL, enrichment among downregulated genes (log_2_FoldChange < -1).

## 3. Methods

### 3.1. Analysis of adhesive and invasive properties of E. coli

*E. coli* isolate ZVL was previously obtained from a patient with Crohn’s disease [3]. *E. coli* cells were cultivated on M9 medium supplemented with 50 mM glucose or 20 mM sodium propionate at 37°C and 180 rpm. The growth curve was monitored using OD at 600 nm and genomic DNA concentration. Cultures in the early logarithmic phase were used for experiments. Adhesive-invasive potential was checked on Caco-2 cells as described earlier [14], with some modifications. The layer of Caco-2 cells was infected by the isolate ZVL at 1:10 ratio. Prior to infection *E. coli* cells were diluted by sterile PBS buffer to 0.1 OD600 centrifuged at 5000 g for 5 min, washed again in PBS, then centrifuged and resuspended in 1 ml of DMEM. The infected cells were incubated for 3 hours at 37°C and 5% CO2. After incubation a subset of the wells was washed two times by PBS to remove non-attached bacteria. Then 0.5 ml of 0.5% Triton X-100 in PBS were added to the wells and incubated for 5 min. The lysate was collected into 0.5 ml of LB broth. These samples contained cells that were attached to or engulfed in Caco-2 cells. The rest of the wells were incubated for 2 hours more. After that the medium was removed and fresh aliquots of 1 ml of DMEM supplied with 300 µg/ml of gentamicin were added and the cells were incubated for 1 hour at 37°C and 5% CO2. After that the wells were washed with PBS and the Caco-2 cells were lysed by Triton X-100 as described above. The obtained samples of *E. coli* in Caco-2 lysates were plated on solid LB agar and cultivated overnight at 37°C. Then the numbers of CFUs were calculated and the values of percent adhesion-invasion and invasion compared to the initial inoculum were determined. The analysis has been carried out in three replicates.

### 3.2. CRISPR-mutants

Plasmids pCas and pTargetF were gifts from Sheng Yang (Addgene plasmids #62225 and #62226, respectively). Knockout of the *exuR* gene was performed by genome editing of *E. coli* using the CRISPR-Cas9 system developed by Jiang et al. [15]. The protospacer sequence was selected using the CHOPCHOP online tool [16]. First, three DNA fragments with overlapping ends were generated by PCR. One of these fragments, encoding the synthetic guide RNA (sgRNA) sequence, was obtained using the exuR-sg4 and sgRNA-R primers and the pTargetF plasmid as a template. Two other fragments encoding homology arms flanking the 500-bp deletion in the *exuR* gene were obtained using *E. coli* MG1655 DNA as a template with primer pairs exuR-bR/exuR-bF and exuR-aR/exuR-aF, respectively. The three resulting DNA fragments were fused into a single chimeric fragment by PCR using primers exuR-sg4 and exuR-aF. This chimeric DNA fragment was then cloned into the pTargetF plasmid via the SpeI and HindIII restriction sites. The nucleotide sequence of the assembled plasmid, designated pTargetT-exuR, was confirmed by Sanger sequencing using primers Targ-scF, Targ-scR, exuR-bF, and exuR-bR. Next the *E. coli* strain ZVL was sequentially transformed with plasmid pCas (conferring kanamycin resistance) and plasmid pTargetT-exuR (conferring spectinomycin resistance) by electroporation. During transformation of bacterial cells with the pTargetT-exuR plasmid, L-arabinose was added to the medium at a final concentration of 10 mM to activate the lambda Red recombination system. *E. coli* ZVL clones with the *exuR* gene deletion were selected on LB agar plates containing kanamycin (50 µg/mL), spectinomycin (50 µg/mL), and L-arabinose (10 mM). Cells were incubated at +30°C throughout all stages of genome editing. Screening of deletion mutants was performed by PCR using primers exuR-scF and exuR-scR. The deletion was then confirmed by Sanger sequencing using the same primers and additionally verified by whole-genome sequencing of the mutant.

### 3.3. RNA Library Preparation and Sequencing

Total RNA was isolated from 1 ml of *E. coli* cultures (CD isolates) using formamide and proteinase K, after chloroform extraction RNA was precipitated with isopropanol. The isolated RNA samples were subjected to DNAse I (Vazyme) treatment for removal of DNA contamination. RNA content was measured using a Qubit RNA HS Assay Kit (Thermo) with a Qubit 2.0 fluorometer (Thermo), purity and quality of RNAs was checked by Nanodrop measurement and agarose gel electrophoresis.

The ribosomal RNA fraction was depleted by hybridization of RNA samples with custom probes (DNA antisense oligonucleotides) followed by RNAse H treatment, DNAse I digestion of probes excess and magnetic beads purification. After chemical fragmentation of depleted purified RNA samples using Mg2+ cDNA synthesis was performed by Magnus reverse transcriptase (“Evrogen”), second strand was synthesised by Klenow fragment (“SybEnzyme”) and RNase H (“NEB”). Then, the resulting double-stranded DNA fragments were purified with magnetic beads and subjected to library preparation.

Library preparation was performed using the MGIEasy Universal DNA Library Prep Set (MGI) following the manufacturer’s protocol. After purification of ligated DNA products an additional size selection step was performed using protocol for samples with 250 bp inserts. Final library concentrations were quantified using a Qubit fluorometer with the Equalbit 1× dsDNA HS Assay Kit (Vazyme). The resulting libraries were converted to DNA nanoballs using the DNBSEQ-G400RS High-throughput Sequencing Kit and sequenced on a DNBSEQ-G400 instrument (MGI) using paired-end 100 bp (PE100) reads, generating approximately 24 million reads per sample.

### 3.4. RNA-seq analysis

Processing of the sequence data was performed using the Nextflow pipeline nf-core/rnaseq (v3.17.0) [9] with the profile ‘singularity’. The complete genome assembly and GTF annotation file of the ZVL strain were used as reference files. The graphics and statistics are taken from the MultiQC report generated by the pipeline. All downstream analysis was performed in R. Estimated counts from Salmon quant.sf files were imported into DESeq2 version 1.38.3 [10] using tximport version 1.26.1 [17]. Differential expression was assessed for two contrasts. The first compared the *exuR* deletion mutant with wild-type ZVL during growth on propionate. The second compared wild-type ZVL grown on propionate with wild-type ZVL grown on glucose. Genes were classified as differentially expressed if the adjusted p-value was < 0.01 and the absolute log_2_FoldChange was > 1.

For Gene Ontology (GO) enrichment analysis, upregulated genes (log_2_FoldChange > 1) and downregulated genes (log_2_FoldChange < −1) were analyzed separately. GO identifiers and their descriptions were extracted from the gene annotation generated by Bakta. GO terms were assigned to their respective categories using GO.db, and only Biological Process terms were retained. Over-representation was tested using the upper tail of the hypergeometric distribution with stats::phyper(). For each term, the test evaluated the probability of observing at least as many annotated genes as were found in the selected set, given the size of the selected set and the frequency of the term in the background. p-values were adjusted across all tested terms using the Benjamini-Hochberg procedure (stats::p.adjust(), method “BH”), separately for each contrast and direction of expression change. Terms with a false discovery rate (FDR) < 0.05 were considered significantly enriched. All significant terms were visualized using ggplot2, and the panels were assembled using patchwork.

## 4. Limitations

A limitation of this study is its focus on a single AIEC strain. Given the genetic diversity of AIEC, the transcriptional and phenotypic patterns observed in ZVL and its *exuR* deletion derivative may not be fully generalizable to other AIEC strains. Differences in gene content and regulatory networks may influence responses to growth medium and *exuR* deletion, comparisons across additional strains would therefore be needed to assess how broadly these findings apply.

## Author Contributions

Investigation, methodology, O.V.; investigation, N.T., D.S., M.G., D.U., A.A., N.B. and A.S.; data curation, writing—original draft preparation, M.M.; conceptualization, methodology, supervision, A.G. All authors have read and agreed to the published version of the manuscript.

## Data Availability Statement

The data are available as a BioProject hosted at NCBI. Data identification number: PRJNA1542820.

## Abbreviations

The following abbreviations are used in this manuscript:

AIEC: Adherent-Invasive *Escherichia coli*
BH: Benjamini-Hochberg
EHEC: Enterohaemorrhagic *Escherichia coli*
FDR: False Discovery Rate
GO BP: Gene Ontology Biological Process
LEE: Locus of Enterocyte Effacement
PCA: Principal Component Analysis
T3SS: Type III Secretion System
WGS: Whole-Genome Sequencing

## References

[1] D. C. Baumgart and W. J. Sandborn, ‘Crohn’s disease’, The Lancet, vol. 380, no. 9853, pp. 1590–1605, Nov. 2012, doi: 10.1016/S0140-6736(12)60026-9.

[2] N. Barnich et al., ‘Adherent-invasive Escherichia coli in Crohn’s disease: the 25th anniversary’, Gut, vol. 75, no. 2, pp. 411–424, Feb. 2026, doi: 10.1136/gutjnl-2025-335331.

[3] O. V. Pobeguts et al., ‘Propionate Induces Virulent Properties of Crohn’s Disease-Associated Escherichia coli’, Front. Microbiol., vol. 11, Jul. 2020, doi: 10.3389/fmicb.2020.01460.

[4] M. J. Ormsby et al., ‘Propionic Acid Promotes the Virulent Phenotype of Crohn’s Disease-Associated Adherent-Invasive Escherichia coli’, Cell Rep., vol. 30, no. 7, pp. 2297–2305.e5, Feb. 2020, doi: 10.1016/j.celrep.2020.01.078.

[5] T. Rosay, A. G. Jimenez, and V. Sperandio, ‘Glucuronic acid confers colonization advantage to enteric pathogens’, Proc. Natl. Acad. Sci. U. S. A., vol. 121, no. 13, p. e2400226121, doi: 10.1073/pnas.2400226121.

[6] T. A. Bessonova, U. D. Kuznetsova, A. T. Magkaev, M. S. Gelfand, and M. N. Tutukina, ‘Effects of Hexuronic Acids on the Expression of Genes for Structural Flagellar Proteins and Regulators of Biofilm Formation in Escherichia coli K-12 MG1655’, Mol. Biol., vol. 60, no. 4, pp. 580–588, Aug. 2026, doi: 10.1134/S0026893326700111.

[7] A. G. Jimenez, M. Ellermann, W. Abbott, and V. Sperandio, ‘Diet-derived galacturonic-acid regulates virulence and intestinal colonization in enterohemorrhagic E. coli and Citrobacter rodentium.’, Nat. Microbiol., vol. 5, no. 2, pp. 368–378, Feb. 2020, doi: 10.1038/s41564-019-0641-0.

[8] T. Lea, ‘Caco-2 Cell Line’, in The Impact of Food Bioactives on Health: in vitro and ex vivo models, K. Verhoeckx, P. Cotter, I. López-Expósito, C. Kleiveland, T. Lea, A. Mackie, T. Requena, D. Swiatecka, and H. Wichers, Eds, Cham (CH): Springer, 2015. Accessed: Oct. 13, 2026. [Online]. Available: http://www.ncbi.nlm.nih.gov/books/NBK500149/

[9] H. Patel et al., ‘nf-core/rnaseq: nf-core/rnaseq v3.17.0 -Neon Newt’, Oct. 2024, doi: 10.5281/zenodo.13986791.

[10] M. I. Love, W. Huber, and S. Anders, ‘Moderated estimation of fold change and dispersion for RNA-seq data with DESeq2’, Genome Biol., vol. 15, no. 12, p. 550, Dec. 2014, doi: 10.1186/s13059-014-0550-8.

[11] O. Schwengers, L. Jelonek, M. A. Dieckmann, S. Beyvers, J. Blom, and A. Goesmann, ‘Bakta: rapid and standardized annotation of bacterial genomes via alignment-free sequence identification’, Microb. Genomics, vol. 7, no. 11, p. 000685, 2021, doi: 10.1099/mgen.0.000685.

[12] G. Sevrin et al., ‘Adaptation of adherent-invasive E. coli to gut environment: Impact on flagellum expression and bacterial colonization ability’, Gut Microbes, vol. 11, no. 3, pp. 364–380, May 2020, doi: 10.1080/19490976.2017.1421886.

[13] M. Amiri, M. Golchin, M. Jamshidian Mojaver, H. Farzin, and A. Hajizade, ‘Enterobactin: A key player in bacterial iron acquisition and virulence and its implications for vaccine development and antimicrobial strategies’, Virulence, vol. 16, no. 1, p. 2563018, doi: 10.1080/21505594.2025.2563018.

[14] M. J. Olivares-Morales et al., ‘Glucocorticoids Impair Phagocytosis and Inflammatory Response Against Crohn’s Disease-Associated Adherent-Invasive Escherichia coli’, Front. Immunol., vol. 9, May 2018, doi: 10.3389/fimmu.2018.01026.

[15] Y. Jiang, B. Chen, C. Duan, B. Sun, J. Yang, and S. Yang, ‘Multigene Editing in the Escherichia coli Genome via the CRISPR-Cas9 System’, Appl. Environ. Microbiol., vol. 81, no. 7, pp. 2506–2514, Apr. 2015, doi: 10.1128/AEM.04023-14.

[16] K. Labun, T. G. Montague, M. Krause, Y. N. Torres Cleuren, H. Tjeldnes, and E. Valen, ‘CHOPCHOP v3: expanding the CRISPR web toolbox beyond genome editing’, Nucleic Acids Res., vol. 47, no. W1, pp. W171–W174, Jul. 2019, doi: 10.1093/nar/gkz365.

[17] C. Soneson, M. I. Love, and M. D. Robinson, ‘Differential analyses for RNA-seq: transcript-level estimates improve gene-level inferences’, Feb. 29, 2016, F1000Research. doi: 10.12688/f1000research.7563.2.

